# Tracking ongoing chromosomal instability using single-cell whole-genome sequencing

**DOI:** 10.1101/2025.11.26.690781

**Authors:** Barbara Hernando, Blas Chaves-Urbano, Maria Escobar-Rey, Marina Torres, Alice Cadiz, Angel Fernandez-Sanroman, Sara Barrambana, Carmen G. Lechuga, Carmen Guerra, Mariano Barbacid, David Gómez-Sánchez, Francisco Sanchez-Vega, Pedram Razavi, Maria Garcia-Perez, Patricia G. Santamaria, Geoff Macintyre

## Abstract

Chromosomal instability (CIN) generates aneuploid genomes that are characteristic of most cancers. While bulk genome sequencing reveals historical CIN, it lacks the resolution to identify ongoing CIN that actively shapes genome evolution. Here, we present a computational framework that leverages single-cell whole-genome sequencing (scWGS) to identify and quantify ongoing CIN by detecting cell-unique copy number alterations and probabilistically mapping them to known CIN signatures. We validated this framework generating *in vitro* models with four types of induced CIN, correctly identifying the induced-CIN type in each case. When applied to cell lines and organoids with ongoing homologous recombination deficiency, our method showed improved identification of sensitivity to PARP inhibition and platinum-based chemotherapy. Analysing scWGS data from 8 triple-negative breast cancers, we linked ongoing impaired non-homologous end joining to subclonal diversification, a finding further supported in cohorts of 179 unmatched primary and metastatic TNBCs and 39 matched cases. Collectively, our results demonstrate that distinguishing ongoing from historical chromosomal instability uncovers a distinct dimension of tumour evolution, suggesting that effective precision oncology will require integrating measurements of both past genomic scars and active mutational processes.

## Introduction

Chromosomal instability (CIN) describes the set of processes that generate numerical and structural DNA changes that typically operate during tumourigenesis(1–3). The result of CIN is an aneuploid genome with changes in DNA copy number ranging from tens of kilobases to whole chromosomes. High-resolution readouts of tumour genome aneuploidy can be achieved using whole-genome sequencing (WGS). The distinct patterns of copy number aberrations (CNAs) seen in these data can then be used to infer which type of CIN operated on the genome(4). For instance, mitotic defects show a pattern of copy number change that is distinct from replication stress. However, this static readout only provides evidence of historical or past CIN, as it represents the average genome state across the population of tumour cells in the sample.

To date, detection of CIN proper, or ongoing CIN, has only been achieved through functional DNA damage response assays, such as γH2AX or RAD51 foci formation(5–9), or through microscopy based methods which show phenomena such as the formation of anaphase bridges or micronuclei(10,11). However, these assays have limitations. They can only detect a fraction of possible CIN-processes, such as mitotic errors or homologous recombination deficiency, but miss others such as replication stress. Furthermore, while they can indicate the presence of DNA damage, they do not necessarily reveal whether the damage was correctly or incorrectly repaired, and whether it persists across cell divisions(12,13).

The identification of ongoing CIN has important clinical implications. For example, current genomic assays, such as Myriad myChoice(14) and HRDetect(15), have successfully leveraged historical homologous recombination deficiency (HRD) signatures to expand the number of patients eligible for PARP inhibitors beyond those with *BRCA1* or *BRCA2* mutations. However, given that CIN is inherently dynamic and tumours evolve over time, the detection of those patients with ongoing HRD that is not yet detectable in bulk DNA sequencing, may further extend PARP eligibility.

The tumour genome provides a rich substrate in which to detect different types of CIN via CIN signature analysis(4,16). However, as mentioned, standard bulk tumour DNA sequencing only provides a readout of historical CIN. To utilise genome signals to identify different types of ongoing CIN, single-cell WGS (scWGS) is required. The genome alterations unique to each individual cell in these data can be used to infer active, ongoing CIN, whereas the alterations shared across cells indicate historical CIN. Proof-of-principle for the detection of ongoing single nucleotide variant (SNV) mutational processes has been achieved via bulk sequencing of single cell clones(17) or pseudobulk of scWGS clones(18). However, current scWGS quality issues prohibit accurate detection of cell-unique SNVs. In contrast, CNAs are readily detectable at single cell resolution, yet a framework for robust identification of cell-unique events and the ability to map these to specific types of CIN is lacking.

To address these limitations, we present a framework for the identification and quantification of ongoing CIN using scWGS data. Our approach leverages CNAs present only in one cell and CIN signatures to describe the active mutational processes driving CIN. Here, we show that our framework can robustly identify different CIN types induced in a collection of *in vitro* models. We apply our approach to pancreatic and breast cancer samples, revealing relationships between ongoing CIN and drug response, and shifts in mutational processes across clones.

## Results

### A framework to identify ongoing chromosomal instability from single cell WGS data

Our framework for identifying and quantifying ongoing CIN consists of three key steps: (1) the generation of genome-wide single-cell DNA copy number profiles from scWGS data, (2) identification of copy number alterations that are unique to individual cells, and (3) assignment of these unique copy number alterations to CIN signatures (**Figure 1a, Methods**). Collectively, the framework outputs whether ongoing CIN was detected, and a probabilistic readout of the type of ongoing CIN.

**Figure 1.**
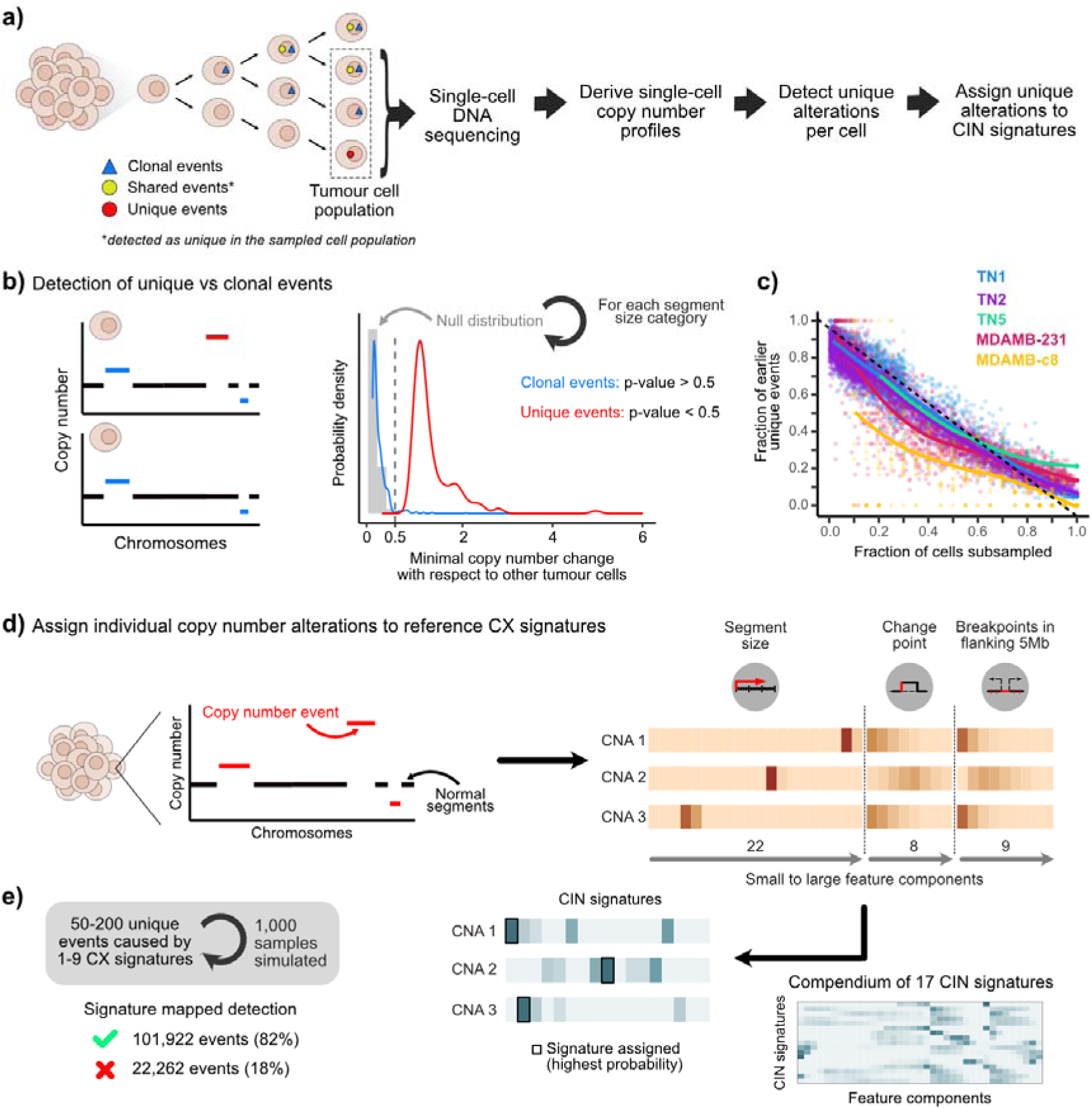
A framework for identifying ongoing chromosomal instability using single-cell whole-genome sequencing data. **a)** Overview of the framework. **b)** Schematic showing our approach to detect cell-unique copy number alterations, including how we determine that the unique events have copy number deviations that are significantly different from a null distribution (see Methods). **c)** Scatter plot showing the fraction of earlier unique events identified (y-axis) when considering only a certain fraction of cells (x-axis) from the whole sample population. Unique events were identified from cell subsets in three triple-negative breast cancer (TNBC) tumours and two TNBC cell lines(19). Earlier unique events were defined as events that appeared unique in the subsampled cell set but were classified as shared when all cells from the population were used. For each sample, cells were randomly selected 20 times. The dashed black line indicates perfect linear correlation. Analyses were performed using samples from Minussi et al, 2021(19) with more than 10 single-cell genome-wide profiles that passed our quality control criteria. **d)** Schematic showing our approach to identify ongoing CIN-related processes. Three copy number features are extracted for each event, which are then probabilistically mapped to the reference CIN signatures. The CIN signature with the highest probability is assigned to the event (see Methods). **e)** In silico validation of our signature mapping approach. Acquisition of copy number alterations caused by a specific signature were simulated for 1,000 samples to evaluate our ability in inferring the underlying signature.

### Single-cell WGS and copy number profiling

The identification of ongoing CIN requires single-cell resolution as only CNAs unique to individual cells (hereafter cell-unique CNAs) provide direct evidence of recent CIN operating on the genome(20–22). Our framework is sequencing platform-agnostic, accommodating data from diverse workflows such as DLP(20), ACT(19) and PicoPlex(23), all of which were used in this study. A critical component of our approach is a rigorous quality control and normalisation pipeline to ensure uniform CNA detection power across all cells. This involves normalising read depths to a consistent target of 15 reads per bin per chromosome copy. Cells which exceed this depth are downsampled accordingly. This equates to approximately 2 million reads for a recommended genome-binning resolution of 100kb. This resolution is crucial for accurate downstream CIN signature assignment. Cells showing evidence of genome replication (S-phase) or noisy profiles are removed from downstream analysis.

### Identification of unique events and ongoing CIN

To identify putative unique events from a collection of single-cell copy number profiles from the same tumour, we first homogenise copy number segment boundaries across cells using a window-based smoothing procedure. This unification of segment boundaries mitigates any unique event misidentification arising from segmentation ambiguities. Copy number is then rounded to the nearest integer and CNAs are classified as clonal if present in ≥90% of cells(19), shared if present in more than one cell, or unique if restricted to a single cell. To ensure robustness of the identified unique events, we apply a test that confirms the observed copy number deviation of the segment from the segments of other cells is significantly different from a null distribution computed from differences in unrounded and rounded segment values across 81,464 segments from 1,441 cells (**Figure 1b**, see **Methods**). We also employed simulations to resolve the evolutionary timing of these ongoing CIN events, acknowledging that the detection of “unique” alterations is inherently dependent on the fraction of cells sampled from the tumour. While unique events in a fully sampled population exclusively represent events from the immediately preceding cell division, sparse sampling captures a mixture of the latest events and earlier events that appear unique due to the absence of sister lineages. Through downsampling analyses, we established a near linear relationship between cell sampling fraction and temporal resolution (**Figure 1c**). Consequently, the percentage of the total population sampled acts as a proxy for the proportion of unique events attributable to the latest cell division, providing an estimate of the temporal recency of the observed ongoing CIN.

### Identification of the type(s) of ongoing CIN

To identify the possible types of ongoing CIN in a sample, cell-unique events are assigned to CIN signatures. To do this, we adapted our previous CIN signature framework(4), redefining the copy number feature embedding space to one which can enable single event assignment, while maintaining the ability to quantify the original CIN signature compendium, all from rounded copy number profiles at 100kb bin resolution (**Figure 1d, Supplementary Note 1**). Using simulations of copy number alternations from known signatures, this approach achieved an accuracy of 82.07% (range across signatures of 67.5-95.7%; **Figure 1e, Supplementary Figure 3**).

For a given sample, not all 17 CIN signatures are likely to be ongoing, as at a bulk level, most samples only exhibit an average of 9 historical CIN signatures with some level of activity(4). However, there could be the possibility that a new signature emerges that is not detected at a bulk level. Therefore, in addition to the bulk signatures, the possible set of ongoing CIN signatures to consider is complemented by the set of signatures detected via assignment of all copy number events across all single cells. Each cell-unique event is then assigned to the CIN signature with the highest probability and the resulting signatures with events assigned can be considered the ongoing CIN types detected.

### Performance assessment using *in vitro* induced-CIN models

To assess the performance of our framework, we induced different types of CIN *in vitro*, performed single-cell DNA sequencing, applied our framework, and then observed whether the ongoing signals detected recapitulated the expected induced CIN type. We created a near-diploid CIN tolerant hTERT-RPE1 *TP53*^-/-^ line, knocked out *BRCA1* or *BRCA2* to induce impaired homologous recombination related CIN; or treated with doxorubicin to induce replication stress related CIN. We also analysed single-clone WGS data(24) from a near-diploid hTERT-HMEC line treated with the spindle assembly checkpoint inhibitor reversine to induce defective mitosis-related CIN. Finally, we performed scWGS on a pancreatic organoid with a known mix of diploid and tetraploid cells, representing the natural occurrence (rather than specific induction) of whole-genome doubling. Collectively, these span the majority of CIN types identifiable via our CIN signature framework.

One caveat to using the cell line *in vitro* systems is that spurious copy number changes can occur in culture attributed to oxidative stress and DNA damage caused by common culture conditions and reagents(25–27). To account for this, we adapted our framework to separate spurious signals (determined in parental cells grown in the same conditions and vehicle-treated) from the induced CIN signals found in the KO/treated lines (**Figure 2a**). The resulting induced CIN signals (encoded in the feature embedding space) were compared with existing CIN signature definitions using cosine similarity to determine the detected ongoing CIN type.

935 unique events from 41 single-cell hTERT-HMEC clones, derived from a parental population treated with reversine(24), showed enrichment of chromosome missegregation-associated signature CX14 (**Figure 2b**), consistent with the CIN type expected to be induced by reversine. 17 unique events identified in 15 hTERT-RPE1 *TP53^-/-^* cells after treatment with doxorubicin matched the replication stress and micronuclei formation related signature CX8 (**Figure 2c**), also consistent with the expected CIN type induced by doxorubicin treatment(28). 16 and 137 unique events detected in 40 and 32 single-cell copy number profiles from hTERT-RPE1 *TP53^-/-^ BRCA1^-/-^* or *BRCA2^-/-^*lines, respectively, matched impaired HR signatures CX2 and CX3 (**Figure 2d**). For the pancreatic organoid system, for which we expect to detect signals of whole-genome doubling (WGD), we detected the subclone-specific unique events by comparing single-cell profiles from the tetraploid clone to the consensus diploid profile, and vice versa. We then applied our mapping procedure to estimate the proportion of unique CNAs putatively caused by a specific CIN signature. As expected, given the organoid had a *BRCA2* mutation, 80% (198) of the unique events in the diploid mapped to impaired HR signatures CX2 and CX3, and 56% (115) in the tetraploid clone. A total of 27 subclone-unique events (13.2%) in the tetraploid clone were assigned to the WGD-linked signature CX4, while an additional 30.4% mapped to the chromosome missegregation-associated signature CX14, consistent with subsequent mitotic errors commonly seen in cells after tetraploidization(29). No unique event in the diploid clone was assigned to CX4, reinforcing the specificity of our approach to capture ongoing mutational processes (Figure 2e).

**Figure 2.**
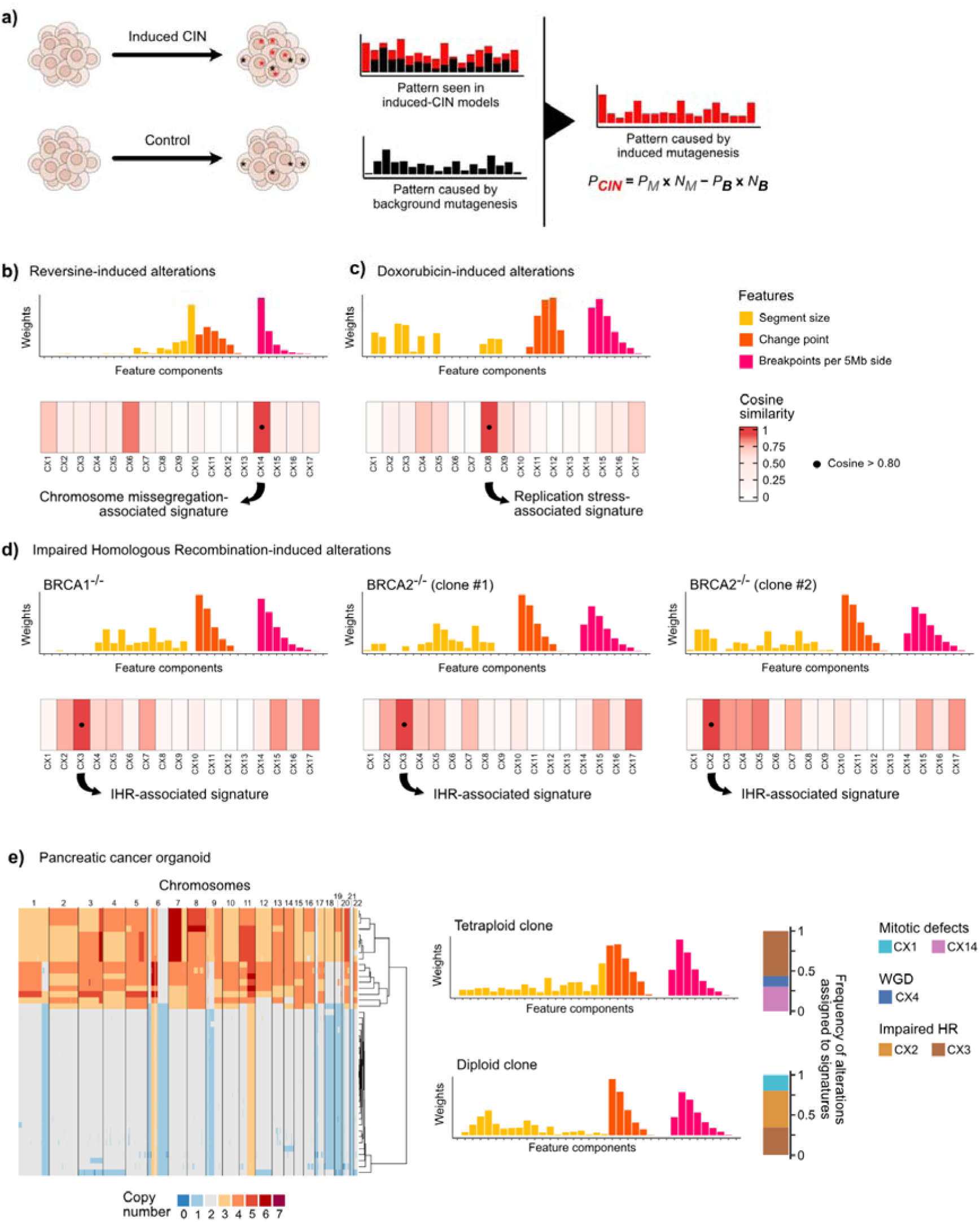
Detection of different types of induced chromosomal instability. **a)** Schematic showing how the genomic scar caused by a CIN-related process is extracted in our in vitro models (see Methods). The CIN-related pattern (P_CIN_) is the result of subtracting the pattern observed in the control (P_B_) to the patterns observed in the induced-CIN model (P_M_). Subtraction is corrected by the mutation rate in both the background (N_B_) and the model (N_M_). **b-d)** Copy number patterns induced by (b) reversine treatment, (c) doxorubicin treatment, and (d) knockout BRCA1/2 genes. Top: Barplots show induced patterns in the 3-feature space. Bottom: Heatmaps show cosine similarities between the induced patterns and the definitions of the reference CX signatures. **e)** Ongoing CIN analysis in a pancreatic cancer organoid with diploid and tetraploid subpopulations of cells. Left: Heatmap showing single-cell copy number alterations. Middle: Barplots show copy number patterns of the subclone-specific unique events. Right: Barplots show frequency of subclone-specific copy number alterations assigned to each CX signature.

### Ongoing CIN can enhance identification of drug sensitivity

Homologous recombination deficiency (HRD) confers sensitivity to treatment with PARP inhibitors(30,31) (PARPi). Patient selection for PARPi, historically based on germline *BRCA1/2* mutations, now includes somatic mutations and signatures of HRD based on genomic patterns of CNAs(14,15,31,32). Here, we hypothesised that these static, bulk-DNA based assays may miss recently acquired HRD, and that identifying these dynamic defects using ongoing CIN could further expand PARPi eligibility. To test this hypothesis, we used the CIN tolerant hTERT-RPE1-Cas9 *TP53*^-/-^ line, knocked out *BRCA2*, sorted and expanded single cell clones for 32 population doublings (PD), performed both bulk and single-cell WGS, and then treated with olaparib or rucaparib to confirm sensitivity to PARP inhibition. We found that the *BRCA2*-deficient model was more sensitive to both PARPi treatments than the wildtype counterpart. We did not detect any HRD-associated copy number alterations in the bulk WGS-derived copy number profiles, indicating that this line would not be considered sensitive to PARP inhibitors at a bulk level, despite the drug treatment showing the contrary (**Figure 3a**). In contrast, unique events detected in the scWGS data showed evidence of impaired HR related CX2 and CX3, indicating sensitivity to PARP inhibitors (**Figure 3a**). These results show that only single-cell, and not bulk WGS, can account for the sensitivity signal of the *BRCA2*-deficient model to PARPi olaparib and rucaparib compared to the *BRCA*-wildtype model.

As impaired HR signatures CX2 and CX3 have also been shown to predict response to platinum-based chemotherapy(4,33), we reasoned ongoing CX2/CX3 might also show enhanced signals over bulk-derived signature predictions. To test this hypothesis, we performed both bulk and scWGS of 4 patient-derived pancreatic adenocarcinoma organoids treated with oxaliplatin. Organoids were classified as either sensitive or resistant based on the previously established prediction rule(33): samples with higher CX3 than CX2 were predicted sensitive, otherwise resistant. Predictions based on ongoing CIN (scWGS) correctly identified the most oxaliplatin-sensitive organoid, which was misclassified as resistant using bulk WGS. In this organoid, 80% of newly acquired events mapped to CX3, indicating a shift from the historical mutational processes predominantly driven by CX2. This provides proof-of-concept that our framework can reveal recent, and potentially current CIN processes associated with treatment responses that are not captured by bulk sequencing (**Figure 3b**). Together, these results provide proof-of-concept that scWGS can reveal ongoing HRD linked to PARPi or platinum sensitivity, that remain hidden in bulk analyses; however, powered cohort studies will be required to fully quantify predictive gains.

**Figure 3.**
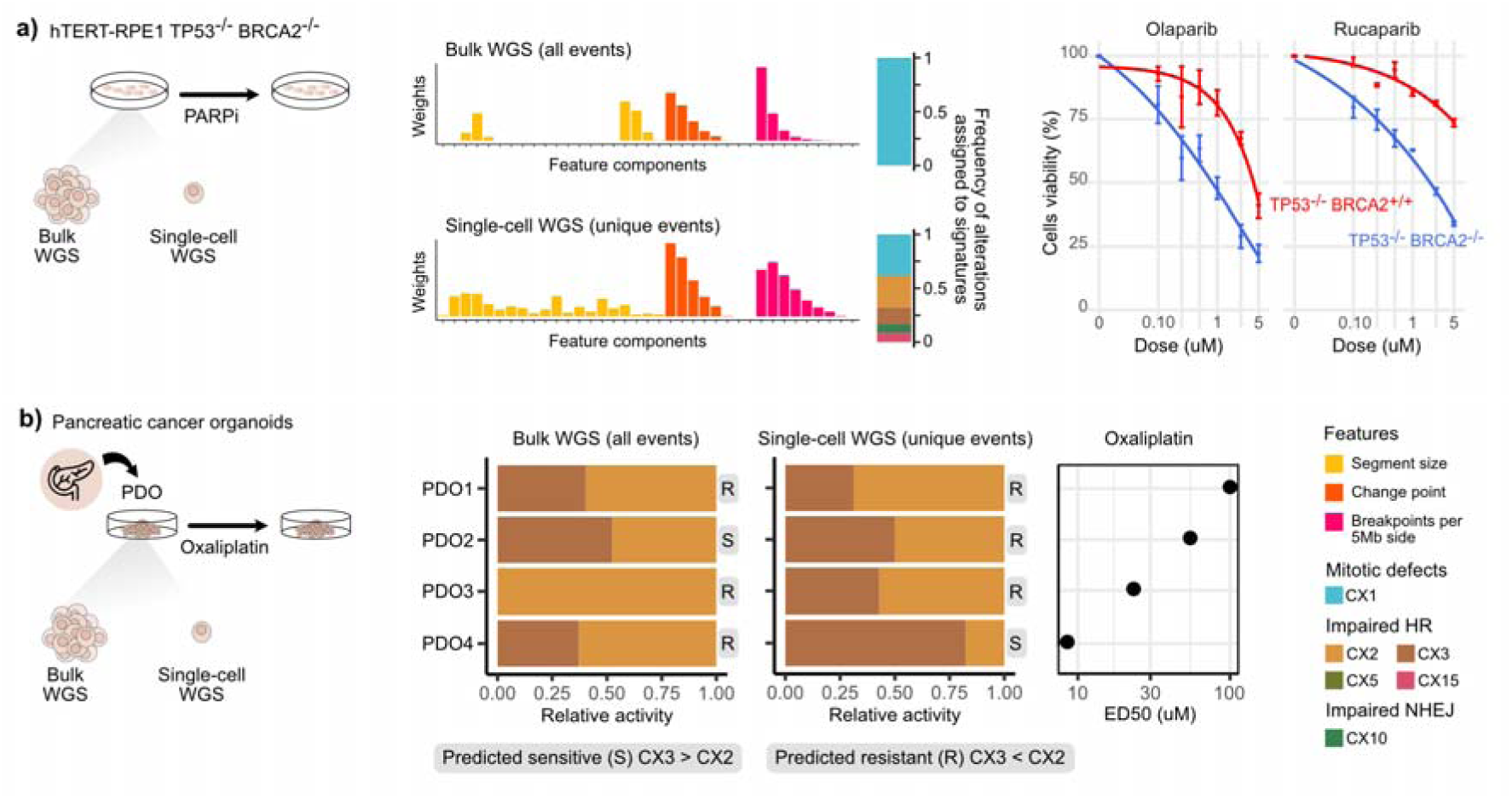
Drug sensitivity prediction based on ongoing chromosomal instability. **a)** Prediction of olaparib and rucaparib response in a BRCA2-deficient model using CX signatures derived from single-cell and bulk whole-genome sequencing. Left: Barplots show copy number patterns of the bulk-detected and cell-unique events. Middle: Barplots show frequency of bulk-detected and cell-unique copy number alterations assigned to each CX signature. Right: dose-response curves for the two PARP inhibitors tested in vitro. A BRCA-wildtype model was used as control. **b)** Prediction of oxaliplatin response in 4 patient-derived pancreatic cancer organoids using CX signatures derived from single-cell and bulk whole-genome sequencing. Organoids were predicted as resistant (R) or sensitive (S) using a previously validated classifier that relies on the CX2/CX3 activity ratio(33). Left: Barplots show CX2/CX3 activity rations derived from bulk WGS. Middle: Barplots show CX2/CX3 activity rations derived from single-cell WGS. Right: ED50 for oxaliplatin.

### Ongoing CIN associates with subclonal diversification in triple-negative breast cancer

Here, we applied our framework to evaluate how ongoing CIN might contribute to the extensive tumour heterogeneity seen in scWGS of a cohort of 8 triple negative breast cancers (TNBC)(19). To achieve sufficient copy number resolution, rather than analyse cell-unique events, we analysed events from subclone pseudobulk copy number profiles. Copy number events shared between all clones constituted historical CIN, and copy number events unique to a subclone were considered to represent ongoing CIN that may have contributed to the subclone diversification.

Clonal *BRCA1/2* inactivation was previously reported in nearly all samples(19). Consistent with this, impaired HR signatures were detected in clonal events and persisted into subclonal epochs (**Figure 4**). WGD-linked signature CX4 was consistently identified as historical in all tumours except in TN5, the only case lacking a pre-diversification WGD event. 7/8 tumours showed the emergence of specific CIN types exclusively in the subclonal epoch. Four patients showed CX10, a signature representing impaired non-homologous endjoining (NHEJ), as ongoing in subclones but absent historically (p-value=0.0008, Binomial test), suggesting NHEJ-deficiency in these tumours may be contributing to subclone diversification. Given that subclonal diversification drives metastasis(34–36), we hypothesized that CX10 would be enriched in advanced disease. We compared CX10 levels across 60 primary(37–39) and 119 metastatic TNBC samples(40) and found significantly higher CX10 levels in the metastatic cohort (p-value=0.0085, Wilcoxon test; **Figure 4d**). Validating this in an additional cohort of 39 paired primary-metastatic TNBC patients(41), we found that – among the 14 patients with detectable CX10 – metastases displayed significantly higher CX10 activity than their matched primaries (p-value=0.025, paired Wilcoxon test; **Figure 4e**). Together, these findings support the association between impaired NHEJ and subclonal diversification in TNBC.

**Figure 4.**
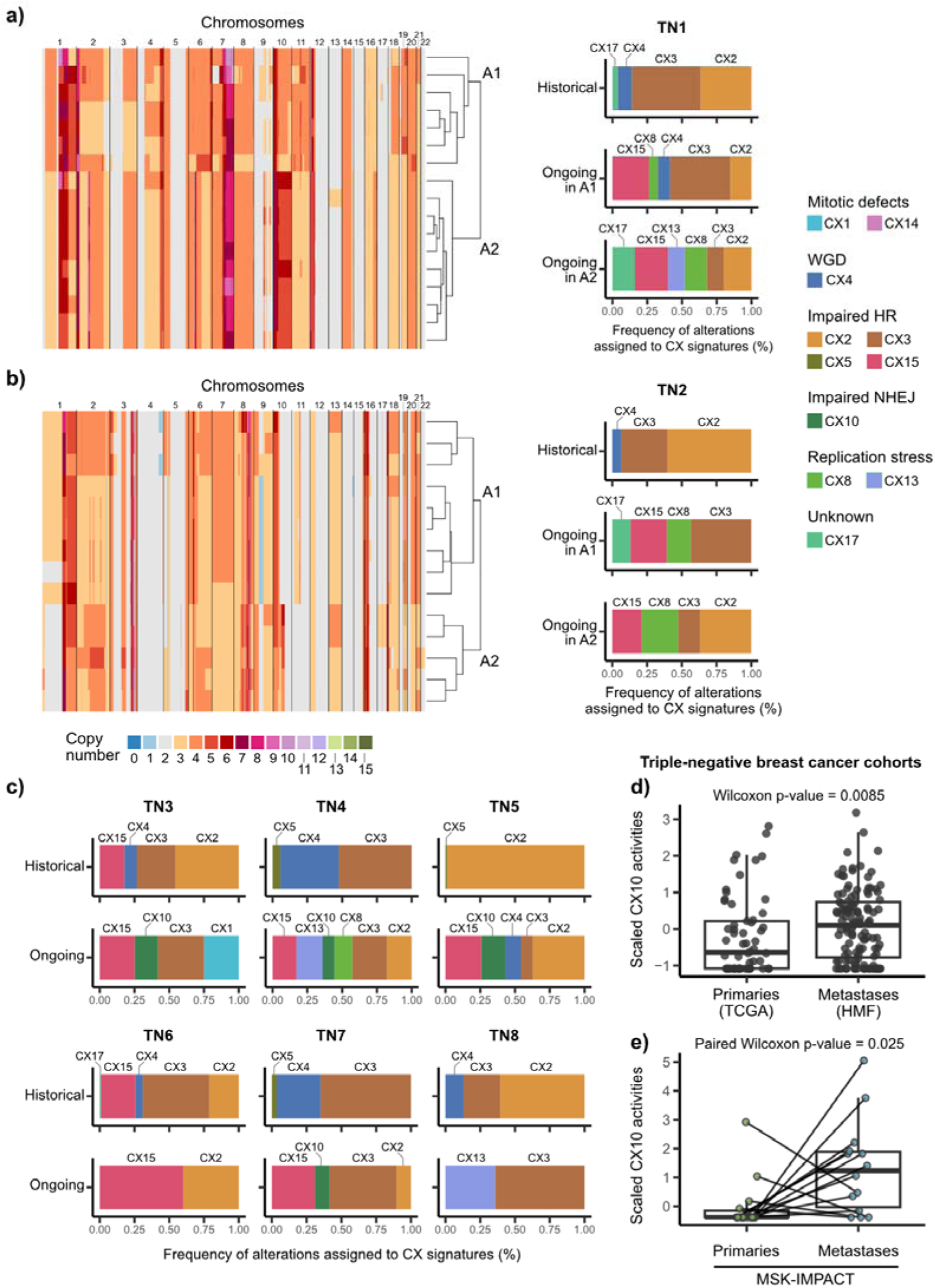
Ongoing chromosomal instability in triple negative breast cancer. **a-c)** Frequency of copy number alterations assigned to historical and ongoing CIN-related processes. Historical CIN was computed using all copy number alterations detected at bulk level, while ongoing CIN was computed using subclone-specific unique copy number alterations (a-b) per branch or (c) per tumour sample. **d)** Activities of the NHEJ-deficient signature CX10 in primary and metastatic triple negative breast cancer samples. Primary samples (n=60) were collected from The Cancer Genome Atlas (TCGA) cohort(37–39), and metastatic samples (n=119) were collected from the Hartwig Medical Foundation (HMF) dataset(40). **e)** Activities of the NHEJ-deficient signature CX10 in primary-metastasis pairs from 39 triple negative breast cancer patients included in the Memorial Sloan Kettering Cancer Center (MSK) dataset. Tumour samples were sequenced using the MSK-IMPACT assay(41).

## Discussion

Here, we present a computational framework for identifying ongoing CIN using single-cell whole genome sequencing, and show that cell-unique CNAs provide a quantitative proxy for recent CIN activity, revealing processes that are invisible to bulk sequencing and may be clinically actionable.

Our method operates in a phylogenetic-tree-free manner, avoiding the need to infer evolutionary trees from copy number alterations, which can be challenging and computationally intensive(42). Most existing approaches for inferring phylogenies unify segment boundaries within clones(19,43,44), potentially altering the true configuration of copy number alterations. Preserving these configurations is essential for accurately tracing the genomic scars left by different mutational processes. By enabling the direct assignment of mutational processes causing cell-unique alterations, our method offers a new tool to investigate the temporal dynamics of CIN-related mechanisms, complementing prior studies that leverage single base substitution signatures(2,45,46).

We demonstrate that the ability to pinpoint the specific CIN mechanisms driving new copy number changes holds promise for improving treatment response predictions. In our analysis, ongoing HRD scars, which were not detected at bulk level, enabled identification of sensitivity to PARP inhibitors in *BRCA2*-deficient models and oxaliplatin in pancreatic organoids. This finding underscores the limitations of using the historical record of genomic alterations for therapy selection and emphasizes the importance of integrating ongoing CIN detection into precision oncology strategies, particularly at early tumour stages. Our analysis of 8 TNBCs(19) also highlighted the importance of ongoing CIN in maintaining tumour evolution and how impaired NHEJ appears to be linked with subclonal diversification. In the original study(19), the authors described that these tumours exhibited a period of early punctuated genomic instability before transitioning to a more moderate rate of ongoing copy number evolution during tumour expansion. Our data are consistent with this being mainly due to ongoing impaired HR. However, subsequent subclonal diversification seems to be associated with the emergence of other types of CIN. Given that all but one patient in the cohort was treatment naive, this represents dynamic, non-treatment related evolutionary pressures. Importantly, we confirmed the association of impaired NHEJ with subclonal diversification in two different TNBC cohorts, including a cohort of patient-matched primary TNBC and metastases.

Our study does, however, have some important limitations. In most practical applications we cannot capture the full cell-unique copy number diversity in a tumour, therefore the “ongoing” CIN identified may in fact be very “recent” and not necessarily active. We also did not perform per-cell-unique-event orthogonal validation (e.g., targeted imaging or breakpoint assays); therefore, while we ensure statistical uniqueness of the events identified, rare segmentation artifacts cannot be fully excluded. Another limitation is that we treat all unique alterations as independent copy number events. Punctuated bursts of copy number alterations have been observed in human cancers(47–50) and model systems(19,51), suggesting that certain alterations may arise as part of the same event that, in some instances, is resolved after several cell cycles. Similarly, WGD triggers a transition from diploid to tetraploid, leading to a genome-wide doubling of copy number alterations (early WGD) often followed by additional gains and losses of genomic regions (late WGD)(52–54).

While our feature encoding successfully detected both early and late WGD *in silico and in vitro*, we have not yet devised an approach to combine CNAs arising from the same punctuated event using scWGS. Another limiting assumption is that copy number alterations remain contiguous with the reference genome, which does not fully represent the complexity of the tumour genome. Further technology developments will be required to enable long-read sequencing or similar at single-cell resolution to overcome many of these limitations.

Ultimately, the ability to quantify ongoing CIN provides a new opportunity for the development of a biomarker that could improve how we select patients for DNA-damage response therapies, augmenting the detection of static genomic scars by revealing the active processes that are driving tumour evolution.

## Acknowledgements

This work was supported by grants to G.M. from the Spanish Ministry of Science and Innovation (Grants PID2019-111356RA-I00 and PID2022-137042OB-I00 funded by MICIU/AEI/10.13039/501100011033 and by FEDER/UE), and by a grant to B.H. from ‘La Caixa’ Foundation (ID 100010434; LCF/BQ/PR23/11980033). The Macintyre and Barbacid groups are hosted by the Centro Nacional de Investigaciones Oncológicas (CNIO), which is supported by the Instituto de Salud Carlos III and recognized as a ‘Severo Ochoa’ Centre of Excellence (ref. CEX2019-000891-S). M.E-R. received the support of a fellowship from the Spanish Ministry of Science and Innovation (grant no. PRE2020-092155 funded by MICIU/AEI/10.13039/501100011033 and by FSE+). B.C-U and A.F-S have received doctoral ‘La Caixa‘ INPhINIT retaining fellowships (IDs 100010434; LCF/BQ/DR23/12000023 and 100010434; LCF/BQ/DR21/11880009). This work was in part supported by grants to M.B. from the CRIS Cancer Foundation, the European Research Council (ERC-AG/695566-THERACAN), the Agencia Estatal de Investigación co-funded with the European Regional Development Fund (ERDF-EU ERDF) (PID2021-124106OB-I00; MCIU/AEI/10.13039/501100011033) and the European Union “NextGenerationEU”/PRTR” (PLEC2022-009255;MCIU/AEI/10.13039/501100011033). M.B. and C.G. are recipients of a CIBERONC Fund (CB21/12/00121). Additional funding included grants to C.G. from the Instituto de Salud Carlos III co-funded by ERDF (PI19/00514), the “Carmen Delgado/Miguel Pérez Mateo Grants of the “Asociación Cancer de Páncreas”. M.B. is the recipient of an Endowed Chair from the AXA Research Foundation. M.G-P. is supported by the Ministerio de Ciencia, Innovación y Universidades (MICIU)/AEI/10.13039/501100011033 and ERDF, EU (Project PID2023-151298OB-I00). This publication and the underlying study have been made possible partly based on data that Hartwig Medical Foundation has made available to the study through the Hartwig Medical Database.

## Author contributions

G.M. and B.H. conceived and designed the study. B.H. developed the methodology and software of the study. B.H., B.C-U., M.E-R., and G.M. contributed to the formal analysis presented in this study. B.H., B.C-U., M.E-R., A.C., A.F-S., M.T., S.B., G.G.L., C.G., M.B., D.G-S., F.S-V., P.R., M.G-P., and P.G.S. provided access to data and/or contributed to gathering, processing and curating data. B.H., P.G.S. and G.M. wrote the manuscript. B.H., P.G.S. and G.M. produced and contributed to the visualisations of the study. G.M. and B.H. supervised the project. All authors had access to all of the data in the study. All authors contributed to the review and the editing of the manuscript. All authors approved the manuscript before submission.

## Methods

### Data reporting and code availability

The source code for reproducing analyses and figures will be available upon publication.

No statistical methods were used to predetermine sample size, and the experiments were not randomised.

All data required for reproducing these analyses will be available upon publication. Raw data generated in this study will be deposited in the European Genome-phenome Archive (EGA) upon publication. The Hartwig Medical Foundation (HMF) clinical, processed and raw sequencing data have restricted access. The Memorial Sloan Kettering Cancer Center (MSK) clinical, processed and raw sequencing data have restricted access. All other data supporting the findings of this study are publicly available without restrictions. Methods Table S1 lists the data used in this study and their respective sources:

**Methods Table S1.**
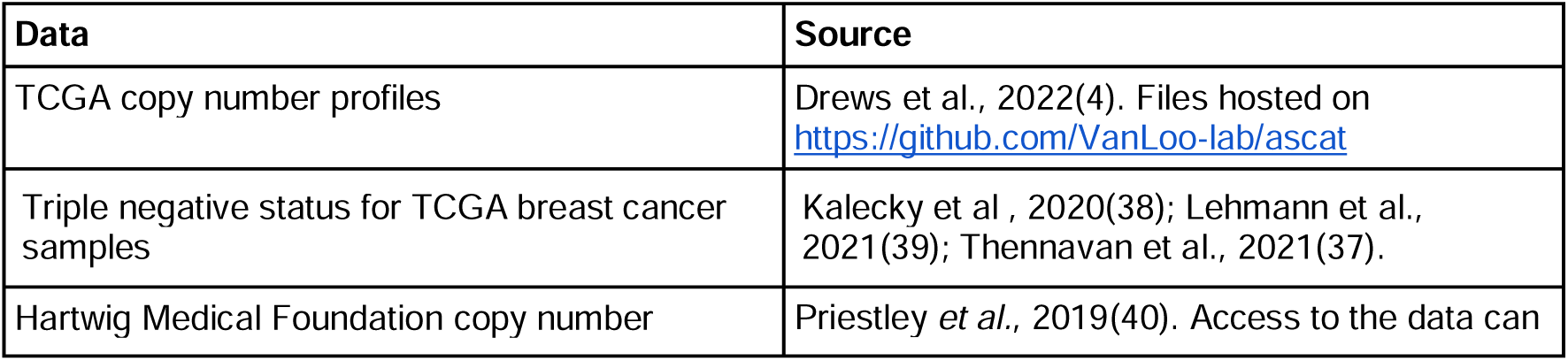
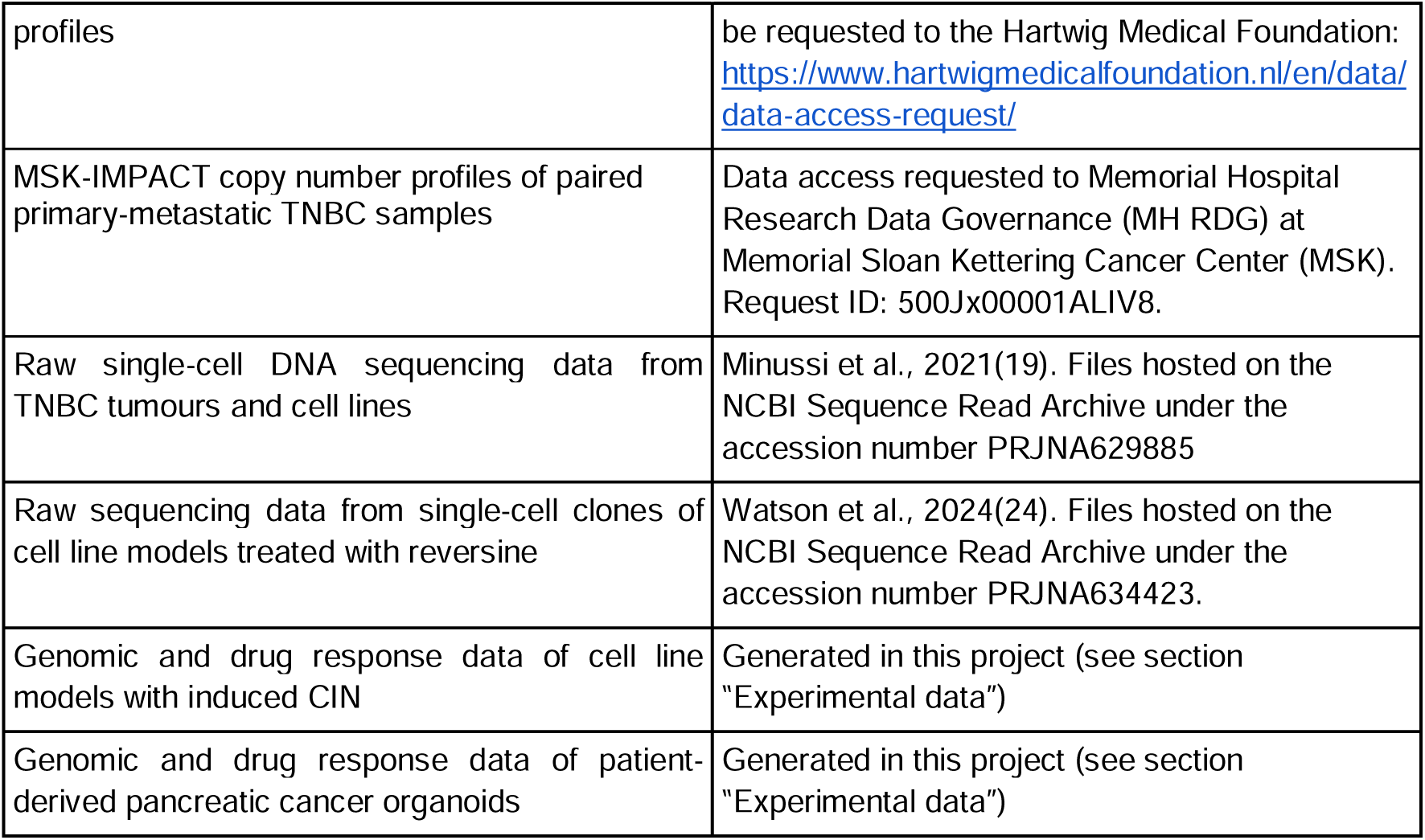
Sources of data used in this study.

### Experimental data

#### Human samples

**Organoid culture.** Five patient-derived pancreatic adenocarcinoma organoids were obtained from patients undergoing surgical resections as first-line treatment at the Hospital Clínico Universitario Virgen de la Arrixaca (Murcia, Spain) under an institutional IRB-approved protocol (CEIC HCUVA-2013/01) and the associated informed consent. Organoids were generated and grown *in vitro* as previously detailed(55). Organoids were recovered from Matrigel (Corning, USA) with TrypLE Express (ThermoFisher) and cells were resuspended in PBS 1X for downstream experiments.

#### Generating cell line models with impaired homologous recombination

**Cell culture.** The hTERT-immortalized normal human retinal pigment epithelial (RPE-1) cell line expressing Cas9 was a kind gift from Dr. Felipe Cortés-Ledesma (CNIO, Madrid, Spain). The hTERT RPE-1 Cas9 *TP53^-/-^*, hTERT RPE-1 Cas9 *TP53^-/-^ BRCA1^-/-^* and hTERT RPE-1 Cas9 *TP53^-/-^ BRCA2^-/-^* cell line models were cultured in Dulbecco’s Modified Eagle Medium Nutrient Mixture F12 (DMEM F12; Sigma Aldrich) supplemented with 10% fetal bovine serum (FBS; Sigma Aldrich), 1% penicillin/streptomycin (Pen/Strep; Solmeglass) and 1% L-glutamine (Gibco). All cell line models were grown at 37°C and 5% CO_2_.

**sgRNA target sequences.** sgRNA targeting the following sequences (3’ to 5’) were used to generate CRISPR knockouts: *TP53* (exon 4) CCATTGTTCAATATCGTCCG; *BRCA1* (exon 12) AAGGGTAGCTGTTAGAAGGC(56); *BRCA2* (exon 11) CTGTCTACCTGACCAATCGA(21). Annealed RNA guides (30 μM) were generated by incubating tracrRNA (5 nmol, Integrated DNA Technologies, IDT), sgRNA targeting each specific locus (100 μM) and nuclease-free Duplex Buffer (Integrated DNA Technologies, IDT) using the following programme in a thermocycler: 95°C (5 min), 70°C (10 min), 45°C (10 min), 20°C (10 min) and 4°C (hold).

**Generation of CRISPR Cas9 knockout cell lines.** hTERT RPE-1 Cas9 or hTERT-RPE1-Cas9 *TP53*^-/-^ cells were seeded in a 60 mm dish and transfected at a 80% of confluence (approx 2.6 x 10^6^ cells) with 2 μL of the corresponding annealed oligo (30 μM) using 17 uL lipofectamine RNAi max (ThermoFisher Cat. #13778075) and Opti-MEM reduced serum medium (Gibco) according to manufacturer’s instructions. Three days after infection, single clones were sorted into 96-well plates on the image based piezo-electric nanoliter dispenser (cellenONE, Scenion) and expanded. hTERT-RPE1-Cas9 *TP53^-/-^*, hTERT-RPE1-Cas9 *TP53^-/-^ BRCA1^-/-^* and hTERT-RPE1-Cas9 *TP53^-/-^ BRCA2^-/-^*clones were selected based on immunoblotting and successful gene editing was confirmed by PCR and DECODR v3.0(57) analysis (https://decodr.org/). The primers (5’ to 3’) used for PCR amplification and Sanger sequencing of targeted *TP53*, *BRCA1* and *BRCA2* loci were: *TP53* forward, GCTGCCCTGGTAGGTTTTCT, TP53 reverse, GAGACCTGTGGGAAGCGAAA; *BRCA1* forward, TCTCAAAGTATTTCATTTTCTTGGTGCC, *BRCA1* reverse, TGAGCAAGGATCATAAAATGTTGG(56); *BRCA2*forward, CCTCAGATGTTATTTTCCAAGCA, *BRCA2* reverse, TCTGCAGAAGTTTCCTCACTAA (**Supplementary Figure 1**).

**Immunoblotting.** Cellular pellet was lysed in buffer [50 mM HEPES pH 7.5, 250 mM NaCl, 5mM EDTA, 1% NP-40, 1mM DTT, 1mM PMSF, 1X protease inhibitor cocktail (Roche) and 1X phosphatase inhibitor cocktail (Roche)] and cells were incubated on ice for 30 min, vortexed every 5 minutes(58). Lysates were centrifuged at 18,000 x g for 1 h at 4°C, and the supernatant was then transferred to a prechilled Eppendorf tube and stored at -80°C. For protein electrophoresis, samples were denatured in 1X NuPAGE™ LDS sample buffer (ThermoFisher) at 70°C during 10 minutes, loaded in an NuPage™ 3-8% Tris-Acetate gel (ThermoFisher), and migrated at 120V for 90 minutes in running buffer (1X NuPage™ Tris-Acetate SDS, ThermoFisher). For transfer, a PVDF membrane (Millipore) was pre-equilibrated in methanol. The proteins were transferred for 4 h at 20V at 4°C. The membrane was blocked in 5% milk in 1X TBS-T at 4°C overnight and then incubated with the respective antibody (see antibodies below) in 5% milk in 1X TBS-T overnight at 4°C. After several washes in TBS-T (3 x 5 minutes), the membrane was incubated for 1 h with the appropriate secondary HRP-antibody at room temperature. After 3 washes in TBS-T, the membrane was developed using Immobilon classico (Millipore) and visualized using a ChemiDoc camera (Bio-Rad). Mouse anti-p53 (1:500, DO-1 sc-126, Cat. #F3023, Santa Cruz Biotechnology), mouse anti-GAPDH (1:1000, NB300-22155, Novus), HRP Goat anti-mouse IgG (1:5000, Cat. #P0447, DAKO) and HRP Goat anti-rabbit IgG (1:5000, Cat. #P0448, DAKO) were used.

**Sanger sequencing**. Genomic DNA from single cell derived clones was extracted using the DNeasy Blood and Tissue Kit (Qiagen, Cat. #69504). PCR amplification of the region of interest was performed using the indicated DNA primers for each gene, PCR products were resolved by agarose gel electrophoresis and PCR product matching expected size were purified using QIAquick Gel Extraction kit (Qiagen, Cat. #280704). Sanger sequencing was performed at the Genomics Unit, CNIO.

#### Bulk low-pass whole-genome sequencing (WGS)

**DNA extraction.** 5 million cells from each cell line model or organoid samples were centrifuged to constitute individual cell pellets that were stored at -80°C until use. DNA extraction from cell pellets was performed using the DNeasy Blood and Tissue Kit (Qiagen, Cat. #69504). Cell pellets were equilibrated at room temperature for 15 minutes to then follow the manufacturer’s instructions. DNA was finally eluted in Tris/HCl (pH 8, 10 mM) and was quantified using the ADNds Quant.iT™ PicoGreen™ kit from Invitrogen (Cat. #P7589).

**DNA library preparation.** Library preparation workflow was executed according to Illumina DNA Prep (Illumina, Cat. #20060060) instructions. IDT® for Illumina® DNA/RNA UD Indexes (Illumina, Cat. #20042666) was used to identify with indexes each sample for sequencing. Quality control of the generated libraries included quantification of the concentration and fragment size assessment of each sample. Library quantification was performed using the ADNds Quant.iT™ PicoGreen™ kit from Invitrogen (Cat. #P7589). Electrophoresis run to assess fragment size was performed with the DNA HiSens Reagent Kit (Revvity, Cat. #CLS760672), which is designed to be used with the HT DNA Extended Range Labchip (Revvity, Cat. #760517). Libraries were pooled together in equal ratios and sequenced by the Genomics Unit of the Spanish National Cancer Research Centre (CNIO) with the NovaSeq™ X Plus system from Illumina, aiming for 10 million reads per fresh frozen sample.

#### Single-cell whole-genome sequencing (scWGS)

**Single-cell isolation.** hTERT RPE-1 Cas9 *TP53^-/-^*, hTERT RPE-1 Cas9 *TP53^-/-^ BRCA1^-/-^*and hTERT RPE-1 Cas9 *TP53^-/-^ BRCA2^-/^* cells were arrested in G1 phase by treating them with 1 μM of Palbociclib for 16 hours. Single cells from cell lines or organoids were single cell sorted using an image based piezo-electric nanoliter dispenser (cellenONE, Scenion) in 96-well PCR plates (Eppendorf Cat. no.: 0030129504). Three to six plates were sorted per model. Cells were selected based on cell line specific parameters of cell diameter, elongation and circularity. Individual cells were dispensed into 96-well PCR plates containing pre-dispensed 1 μL (PicoPLEX) or 300 nL (mDLP+) of 30 mM Tris pH 8, then stored at - 20°C.

**Single-cell DNA library preparation.** Single-cell DNA whole genome sequencing libraries were performed using the SMARTeR PicoPLEX Gold Single DNA-Seq Kit (PicoPlex Gold, Takara) or using a modified version of the Direct Library Preparation Plus protocol previously described (mDLP+)(20).

For the Picoplex protocol, cells (in 1 μL 30 mM Tris pH 8) were subjected to an enzymatic and heat lysis for efficient release of genomic DNA. Then, a pre-amplification step of the genomic material was performed, followed by a clean up step for removing the excess of pre-amplification primers (AMpure XP beads, Beckman Coulter). Finally, an amplification step was conducted for exponential amplification of the DNA libraries and IIlumina-compatible indexing.

For the mDLP+ protocol, cells (in 300 nL 30 mM Tris pH 8) were first subjected to a lysis step. We added 300 nL of freshly prepared 2X protease lysis buffer [800 μL 2X lysis buffer (30 mM Tris pH 8, 1% Tween-20, 1% Triton-X-100)] and 200 μL QIAGEN Protease (1.36 Anson units/mL prepared in nuclease-free water) to each well using the Mantis Liquid Dispenser (Formulatrix), followed by 10 min incubation at 55°C and 15 min at 75°C. DNA tagmentation was performed by adding 1 μL of TD buffer and 0.5 μL of ATM buffer mixture (Nextera XT DNA Library Preparation Kit, Illumina) and incubated at 55°C for 5 min. The tagmentation reaction was neutralised by 5 min incubation at room temperature with 0.5 μL NT buffer (Nextera XT DNA Library Preparation Kit, Illumina). Library amplification was performed by adding 4 μL (2X) Equinox Amplification Master Mix (Equinox Library Amplification Kit, Watchmaker Genomics) and 1,5 μl of 2 μM unique dual indexed adapter pool (Illumina DNA/RNA UD Indexes, Illumina) using the following PCR conditions: 72°C (5 min), 98°C (45s), 14 cycles of 98°C (15s), 60°C (30s), 72°C (45s), followed by 72°C (5 min), with the lid temperature at 105°C. The quality and quantity of the single-cell DNA libraries were assessed with DNAds Quant-iT™ PicoGreen™ kit on FLUOROSKAN (ThermoFisher Scientific) and LabChip DNA High Sensitivity Reagent kit on LabChip GX Touch Nucleic Acid Analyzer (PerkinElmer) according to the supplier’s recommendations. Libraries performed using the PicoPlex protocol with DNA concentration lower than 5 ng/μL were discarded. Pooled libraries in equimolar ratios were purified using 1X AMpure XP (Beckman Coulter) beads and sequenced using Novaseq™ X Plus system from Illumina.

#### Drug treatments

**Genotoxic agents.** hTERT RPE-1 Cas9 *TP53^-/-^* cells were treated with a sublethal dose of doxorubicin (5 nM(59)) (Cat #D1515 Sigma-Aldrich) for 10 PDs, and the drug was refreshed every 3 days. PARP inhibitors olaparib and rucaparib (a gift from the Experimental Therapeutic Programme at the Spanish National Cancer Research Centre (CNIO)) were used as detailed below.

**Drug sensitivity assay.** Cell viability was assessed in hTERT RPE-1 Cas9 *TP53^-/-^* and hTERT RPE-1 Cas9 *TP53^-/-^ BRCA2^-/-^* cells. 1,000 cells per well of each cell line were seeded in opaque 96-well plates (ThermoScientific) and treated at increasing concentrations of olaparib and rucaparib (0.1, 0.25, 0.5, 1, 2.5 and 5 μM) for 6 days. Cell viability was assessed using Cell Titer Glo assay kit (Promega) according to manufacturer’s instructions. Luminescence was read on an EnVision Multimode plate reader 2104 (Perkin Elmer). Measurements were corrected for background absorbance (DMSO), and survival percentages were calculated relative to vehicle-treated (DMSO) control cells.

**Drug treatment of pancreatic cancer organoids.** Viable single cells (4,000) from 4 pancreatic cancer organoids were plated in 4 µl of Matrigel in 96-well tissue culture plates in duplicates as previously described(55). After 72 h organoids were treated with a range of oxaliplatin concentrations (0.003-50 μM) and viability was assayed 96 h later with Cell Titer Glo (Promega). Drug-response curves were fitted using the R package *drc(60)* to estimate ED50 values for each organoid. To ensure consistent curve fitting, the concentration range was extended up to 100 μM for model fitting. If cell viability did not decrease by more than 20%, the ED50 value was assigned to the highest fitted concentration (**Supplementary Figure 2**).

### External data

#### Watson et al. 2024

We downloaded publicly available sequencing data from normal diploid human telomerase reverse transcriptase (hTERT)-immortalized human mammary epithelial cells (hTERT-HMECs) treated with reversine(24), a potent inhibitor of the spindle assembly checkpoint(61). Whole-genome sequencing data from post-treated single-cell clones was processed to derive copy number profiles (see section “Generating copy number profiles”). Clones were derived by allowing an individual HMEC cell to expand for 20 population doublings (PD). For ploidy selection, we used the ploidies determined via propidium iodide staining by the authors.

#### Minussi et al. 2021

We downloaded publicly available single-cell sequencing data from 8 triple-negative breast cancer patients(19). Only cancer cells were sorted and sequenced in this work. Sequencing data was processed to derive single-cell copy number profiles (see section “Generating copy number profiles”). We also derived subclone-specific copy number profiles by concatenating sequencing reads of all cells belonging to the subclone. Ploidy selection was informed by the flow cytometry data reported by the authors, who isolated cell populations of defined ploidy for each tumour sample.

#### Primary and metastatic triple negative breast cancer samples

A total of 60 primary triple-negative breast cancer (TNBC) tumours were compiled from the Tumour Cancer Genome Atlas (TCGA) cohort(37–39). ASCAT-derived TCGA copy number profiles were downloaded from https://github.com/VanLoo-lab/ascat. We used three different datasets to annotate triple-negative status(37–39,62), and only tumours consistently annotated as triple-negative across all three were retained for downstream analysis.

We also analysed 119 metastatic triple-negative breast cancer samples from the Hartwig Medical Foundation (HMF) dataset(40). We downloaded PURPLE-derived copy number profiles from HMF, and processed them as previously described(63). Triple-negative annotations were retrieved directly from the HMF metadata.

For each copy number profile, we computed the genome-wide distributions of the 3 copy number features, and quantified activities of CIN signatures using the linear combination decomposition function from YAPSA(64) as previously described(63). CIN signature activities were finally scaled within primary and metastatic samples to enable comparison across cohorts. Wilcoxon rank-sum test was used to compare CX signature activities between primary and metastatic TNBC tumours.

#### Paired primary-metastatic TNBC samples from MSK-IMPACT

We processed 82 TNBC tumours comprising 39 matched primary-metastatic pairs from the same patients. We used tumour-normal pairs sequenced with the MSK-IMPACT targeted panels(41) to generate genome-wide absolute copy number profiles using FACETS(65). When patients had more than one primary or metastatic sample available, we retained the profile with the highest quality and/or purity. The final dataset therefore consisted of 39 primary tumours and 39 matched metastatic tumours. For each copy number profile, we then quantified CIN signature activities as described in the previous section. Paired Wilcoxon signed-rank test was used to compare CX10 activities between primary TNBC tumours and their matched metastases.

### Generating copy number profiles

Reads were aligned as single-end against the human genome assembly GRCh37 using BWA-MEM (v0.7.17). Duplicate reads were then identified and marked using samtools-markdup (v1.15). After alignment, absolute copy number profiles were fitted from sequencing data generated across various high-throughput technologies.

We used the QDNAseq R package(66) to count reads within 100 kb bins. Bins mapped to centromeres and regions of undefined sequence in the reference genome hg19 were excluded. Sex chromosomes were also excluded for generating copy number profiles. Read counts were initially corrected for the relationship between sequence mappability and GC content. After segmentation, absolute copy numbers were inferred across a ploidy range (1.5 < ploidy < 8) and a purity fixed at 1, as previously described(33). If the sample’s ploidy was known, a narrower range (±0.5 of the ploidy) was used. For consistency, samples were downsampled to 15 reads per bin per chromosome copy, and samples not meeting this sequencing depth were excluded from downstream analyses.

The optimal mathematical solution was considered the one with the lowest root mean squared deviation (RMSD) between the non-rounded and the rounded copy numbers for all bins. Fitting solutions were manually reviewed and adjusted if necessary.

Following absolute copy number fitting, samples were rated using a star system as previously described(16,33), and those with 1-star were discarded for downstream analyses. For processing single-cell WGS data, we excluded cells with excessive noise, defined as having a standard deviation of bin counts higher than the median across all cells. We also excluded cells with fragmented profiles, characterized by having breakpoint counts exceeding two standard deviations above the mean across all non-noisy cells. Finally, we rounded the copy number values of each segment to the nearest integer and merged adjacent segments with the same copy number to generate the copy number profile for each cell.

### Identifying cell-unique copy number alterations

We classified copy number alterations detected in single cells into three categories: clonal, shared and unique. Unique copy number alterations were defined as segments present in a cell with a distinct copy number value compared to all other cells. Shared copy number alterations were defined as segments with the same copy number value in more than one cell, while clonal alterations were those found in at least 90% of all cells(19).

Further details of the heuristic approach applied to ensure robust classification of unique copy number alterations will be available upon publication.

### Mapping signatures to individual copy number segments

We developed a probabilistic approach to assign individual copy number events to CIN signatures. To enable single-event mapping resolution, we adapted our previous CIN signature framework(4) by redefining the copy number feature space, allowing the analysis of alteration patterns per *individual event* rather than per *genomic region*. The full details of the development and *in silico* validation of the redefined encoding framework are outlined in **Supplementary Note 1**, which will be publicly available upon publication.

